# Population genomics resolves cryptic species of the ecologically flexible genus *Laspinema* (cyanobacteria)^1^

**DOI:** 10.1101/2024.02.26.582125

**Authors:** Petr Dvořák, Svatopluk Skoupý, Hana Jarošová, Kateřina Páleníčková, Aleksandar Stanojković

**Affiliations:** Palacký University Olomouc, Faculty of Sciences, Department of Botany, Šlechtitelů 27, 78371 Olomouc, Czech Republic

**Keywords:** cyanobacteria, cryptic species, evolution, gene flow, new species, phylogenomics, population genomics

## Abstract

Cyanobacterial taxonomy is entering the genomic era, but only a few taxonomic studies have employed population genomics, which provides a framework and a multitude of tools to understand species boundaries. Phylogenomic and population genomic analyses previously suggested that several cryptic lineages emerged within the genus *Laspinema*. Here, we apply population genomics to define boundaries between these lineages and propose new cryptic species, *Laspinema olomoucense* and *Laspinema palackyanum*. Moreover, we sampled soil and puddles across Central Europe and sequenced the 16S rRNA and 16S-23S ITS of the isolated *Laspinema* strains. Together with database mining of 16S rRNA sequences, we found that the genus *Laspinema* has a cosmopolitan distribution and inhabits a wide variety of habitats, including freshwater, saline water, mangroves, soil crusts, soils, puddles, and the human body.

## Introduction

Cyanobacteria exhibit great phenotypic variability, which enabled a description of several thousand species (Nabout et al. 2013, Dvorák et al. 2017). However, soon after the introduction of DNA sequencing and phylogenetic analyses, it became obvious that phenotype evolution often does not follow genome evolution (Turner et al. 1999). The two main patterns emerged. First, the evolution of cyanobacteria is muddled with many events of parallel and convergent morphological evolution. The most prominent example of all is hidden under the name *Synechococcus*. These simple rod like cells are cosmopolitan in diverse environments and belong to the most abundant organisms on the planet (Flombaum et al. 2013). *Synechococcus*-like cyanobacteria have evolved independently at least 12 times over three billion years (Dvořák et al. 2014a). Some of these polyphyletic lineages have already been transferred into the new genera (Dvořák et al. 2014b, Komárek et al. 2020). This phenomenon is widely spread across most of the cyanobacterial lineages and it has led to the boom of more than a hundred of new genera within the last decade (Komárek et al. 2014, Dvořák et al. 2015, 2021). Second, the new monophyletic genera defined based on the 16S rRNA phylogeny are often composed of several cryptic species, i.e., no morphological features distinguishing them. For instance, the genus *Oculatella* is composed of seven species, which are recognized only using the 16S rRNA and 16S-23S internal transcribed spacer (ITS) secondary structures (Osorio-Santos et al. 2014).

The phylogenetic reconstruction based on the 16S rRNA certainly provided a powerful tool in taxonomic research and most of the new species and genera are identified using 16S rRNA sequence. However, the evidence is growing that 16S rRNA alone cannot distinguish species units in many bacterial phyla (Fox et al. 1992, Hassler et al. 2022) To facilitate this problem, it became a common practice among the taxonomists of cyanobacteria to employ 16S-23S ITS secondary structures since it was introduced by Boyer et al. (2001) because this locus is more variable than 16S rRNA. However, while the inference of ITS secondary structures could be effective in many taxa, its diversity may be too low in others. The most recent example represents the cosmopolitan genus *Argonema*. The ITS secondary structures were the same among the two species *A. galeatum* and *A. antarcticum* (Skoupý et al. 2022). The two species were recognized based on the phylogeny of 16S rRNA, morphological apomorphy, and the genome difference. Moreover, 16S rRNA and ITS operon could be present in several copies within the cyanobacterial chromosome. These copies often differ more within the chromosome of a particular strain than between the species or even genera (Johansen et al. 2017).

The number of cryptic species is possibly much larger because the 16S rRNA widely underestimates the genome-wide diversity (Hassler et al. 2022). However, taxonomic studies investigating the genome sequences among the cyanobacterial species are only scarce, although whole genome sequencing became more affordable within the last decade with 5255 genome assemblies of cyanobacteria submitted to the GenBank (https://www.ncbi.nlm.nih.gov/, accessed 23^rd^ February 2024). For example, the genome sequences were employed to an erection of the new genera *Moorena* (Engene et al. 2012), *Elainella* (Jahodářová et al. 2018), and *Argonema* (Skoupý et al. 2022). If the whole genome sequence is present in the taxonomic study, each taxon is usually represented by one genome sequence.

Here, we will dive deeper into the genome diversity of species using population genomics. We recently proposed a framework for the application of population genomics in cyanobacterial taxonomy (Dvořák et al. 2023). We argued that the wealth of the sequencing data enables detailed inference of the evolutionary patterns within the populations and species of cyanobacteria. We could reconstruct the drivers of the speciation and define the boundaries between the species more precisely and closer to nature. Thus, the delimitation is not hurdled by the resolution of conservative markers or low morphological diversity (as we showed above). Population genomics represents the most powerful tool for understanding and delimitating of cryptic species (Skoupý et al. 2024, Stanojković et al. 2024).

The genus *Laspinema* was derived from *Phormidium* based on two *Laspinema* strains isolated a thermal spring with elevated radiation in Iran (Heidari et al. 2018). Besides the type species *L. thermale*, the genus contained another species: *L. etoshii* (previously *Phormidium etoshii*; (Dadheech et al. 2013)), *L. lumbricale* and *L. acuminatum* (previously “*Oscillatoria acuminata”* PCC 6304) (Heidari et al. 2018, Zimba et al. 2021). Most recently, we sequenced the whole genomes of 8 *Laspinema* strains from a small puddle in the city of Olomouc (Czechia). The strains were first identified as *Laspinema* based on the morphology and 16S rRNA. However, the phylogenomic and population genomics analyses showed that the *Laspinema* strains exhibit extensive cryptic diversity and belong to at least two cryptic lineages. The estimated recombination rate was low between the species and higher within the species, suggesting a barrier to gene flow between the highly diverged lineages that remain cryptic. We identified genes associated with nutrient uptake and response to environmental stimuli like light intensity or desiccation, which may have been involved in the divergence via ecological selection (Stanojković et al. 2022).

Here, we will explore the cryptic diversity within the species of *Laspinema*. We will employ population level sampling and genome sequences to propose new cryptic species. Furthermore, we will infer the distribution within the *Laspinema* utilizing the database sequencing data and our sampling effort in Central Europe.

## Material and methods

### Strain maintenance and morphological characterization

Strains with the genome sequence were adopted from Stanojković et al. (2022) and new samples were collected at several localities in Czechia and Poland (complete list of strains and localities; Table S1). The strain isolation was performed as in our previous taxonomic proposals (Dvořák et al. 2017, Skoupý et al. 2022). The isolated strains were maintained under the following conditions: Zehnder medium, temperature 22±1°C day temperature, 18±1°C night temperature, illumination 20 µmol photons m^−2^.s^−1^, light regime 16h light/8h dark. We characterized the morphology of strains using a Zeiss AxioImager microscope with high resolution camera AxioCam HRc 13MPx. We focused on the cell shape, dimensions, filament apices shapes, reproduction, and color. We selected seven strains with available genomes to assess the morphological differences between the cryptic species. Fifty widths and lengths of cells for each strain of both species were measured under 1000x magnification, and differences between the cell dimensions were assessed using a one-way ANOVA test in R (R Core Team 2021).

### PCR amplification and sequencing

Genomic DNA was extracted from 50 mg of fresh biomass using the DNeasy UltraClean Microbial Kit (Qiagen, Hilden, Germany), following the manufacturer’s instructions. The quality of the extracted DNA was assessed by agarose gel electrophoresis (1.5% agarose gel, GelRed; Biotium, California, USA). The DNA concentration was quantified using a NanoDrop 1000 spectrophotometer (Thermo Fisher Scientific, Wilmington, Delaware, USA). A partial 16S rRNA gene sequence and a whole 16S-23S ITS sequence were amplified by PCR using the primers P2 (5’- GGGGAATTTCCGCAATGGG-3’) and P1 (5’-CTCTGTGTGCCAGGTATCC-3’). The PCR reaction was performed in a total volume of 40 μL containing 17 μL of sterile RNase-free water, 20 μL of EmeraldAMP Master Mix (Takara Bio Europe SAS, Saint Germain en Laye, France), 1 μL of each primer, and 1 μL of template DNA (50 ng/μL). The amplification conditions were as described by Skoupý et al. (2022). The PCR products were purified using the E.Z.N.A Cycle Pure Kit (Omega Bio-Tek, Georgia, USA) according to the manufacturer’s instructions. The PCR products were sequenced using the Sanger method (Macrogen Europe B.V., The Netherlands) using two additional primers, P5 (5’-TGTACACACCGCCCGTG-3’) and P8 (5’-AAGGAGGTGATCCAGCCACA-3’). The acquired 16S rRNA and 16S-23S ITS sequences were trimmed and assembled using Sequencher 5.0 software (Gene Codes Corporation, Ann Arbor, MI, USA). The sequences were also uploaded to the NCBI database (accessions, Table S1).

### Phylogenetic, phylogenomic reconstructions and ITS secondary structures

The whole-genome phylogeny of the whole cyanobacterial phylum was reconstructed based on representatives of cyanobacterial clades, all sequenced genomes of *Phormidium* and *Oscillatoria* available in NCBI database (https://www.ncbi.nlm.nih.gov/), and *Laspinema* genomes from Stanojković et al. (2022) (full list; Table S2). The multiple sequence alignment (MSA) was produced by OrthoFinder 2.3.1 (Emms and Kelly 2015), which searched orthologs of investigated taxa with default settings. The maximum likelihood (ML) phylogenetic tree was reconstructed in IQ-TREE 1.6.5 (Nguyen et al. 2015). The best model was selected using Modeltest implemented in IQ-TREE (Kalyaanamoorthy et al. 2017) based on the Bayesian information criterion (BIC) as follows – LG+F+G4. The tree topology was tested by 2000 ultrafast bootstrap re-samplings (Hoang et al. 2018).

The phylogenomic tree of the *Laspinema* species was inferred based on the complete proteoms as described above. *Ancylothrix* sp. D3o was selected as the outgroup based on the phylogeny of the whole cyanobacterial phylum. The model for the ML reconstruction was identified again as LG+F+G4.

Next, we used all sequences of 16S rRNA and 16S-23S ITS found in NCBI and our newly sequenced strains. We searched the NCBI using BLAST (https://blast.ncbi.nlm.nih.gov/Blast.cgi) to identify the 16S rRNA and 16S-23S ITS sequences tentatively belonging to *Laspinema* using a sequence of *L. palackyanum* D2a as a query.

Finally, we searched NCBI using BLAST for all 16S rRNA sequences of *Laspinema*. All sequences were added to the 16S rRNA phylogenetic dataset from Skoupý et al. (2022). This way, we could identify the monophyletic clade of *Laspinema* among all cyanobacteria using two datasets – 16S rRNA only and as well as concatenated 16S rRNA and 16S-23S ITS. The MSA were aligned in Aliview (Larsson 2014) using the Muscle 5.1 (Edgar 2004) with the default settings. The alignment was trimmed using trimal 1.4 (Capella-Gutiérrez et al. 2009) with settings-automated1. We used IQ-TREE again with the same settings. TIM2+F+I+G4 (16S rRNA and 16S-23S ITS) and GTR+F+I+G4 (16S rRNA only) were identified as the best models for the ML reconstruction. The 16S rRNA and 16S-23S ITS dataset was further analyzed using Bayesian inference in MrBayes 3.2.3 (Ronquist et al. 2012) with two separate runs (each with 3 heated and 1 cold chain) for 4 million generations and GTR+I+G model. The sampling frequency was each 1000th generation and 25% of trees were discarded as burn–in. The consensus tree was constructed using a 50% majority rule. PRSF values reached 1 and ESS values were all above 1700. Maximum parsimony inference for 16S rRNA and 16S-23S ITS dataset was performed in MEGA X software (Kumar et al. 2018) using the subtree pruning regrafting search method. All trees were edited in FigTree 1.4.4 (https://github.com/rambaut/figtree/releases) and Inkscape (https://inkscape.org/).

We estimated secondary structures of D1-D1’ and Box B helices using Mfold RNA folding form with default options except for structure draw mode, which was changed to ‘untangle with loop fix’ (Zuker 2003). The regions of ITS of D1-D1’ and Box B helices were identified using CIMS at https://www.phylo.dev/ (Labrada et al. 2023).

### Gene and genome similarity estimates

The similarity matrix of 16S rRNA sequences was estimated using p-distance based on the MSA described above. The analysis was performed in MEGA X software (Kumar et al. 2018) using uniform rates, and composite likelihood, while the gaps were deleted pairwise. The average nucleotide identity (ANI) was calculated using fastANI with the default settings (Jain et al. 2018).

### Taxonomy and nomenclature

The new species was proposed using a combination of biological species concept adapted to cyanobacteria (Dvořák et al. 2023) and monophyletic species concept sensu Johansen and Casamatta (2005). The species is proposed according to the rules of the International Code of Nomenclature for Algae, Fungi, and Plants (https://www.iapt-taxon.org/nomen/main.php).

## Results

### The phylogeny of Laspinema

We reconstructed the phylogenomic tree of the whole phylum cyanobacteria based on the MSA composed of ∼165k amino acid positions. The major tree branching resembled published phylogenies (Jahodářová et al. 2018, Skoupý et al. 2022). The genomes of *Laspinema* formed a well-supported monophyletic clade sister to *Oscillatoria* sp. SIO1A7, *Ancylothrix* sp. D3o, *Baalinema simplex*, and a clade of putative *Phormidium* species (Figure 1). Then, we inferred the whole-genome phylogeny within the genus *Laspinema* based on ∼587k amino acid MSA (1723 single-copy orthologs). The phylogeny revealed several clades. *Laspinema palackyanum* formed a separate cluster from “*Oscillatoria acuminata”* PCC 6304, “*Phormidium pseudopriestleyi”* FRX01, *Laspinema* sp. D2d and *L. olomoucense* (Figure 2). The clades of *L. palackyanum* and *L. olomoucense* were significantly supported.

**FIGURE 1.**
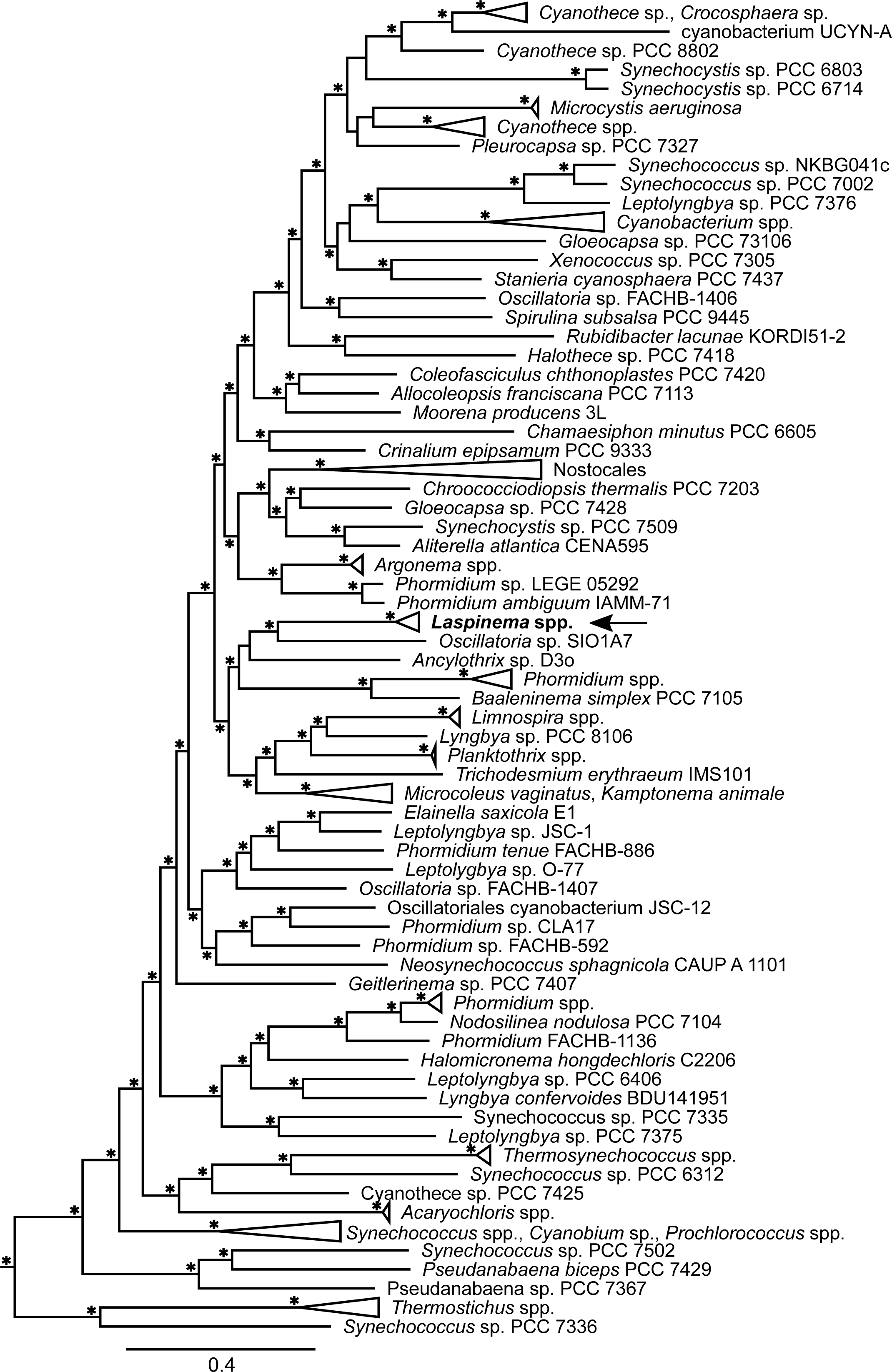
The whole-genome phylogeny of cyanobacteria. The maximum likelihood was reconstructed on the amino acid multiple sequence alignment and rooted to the *Gloeobacter* spp. (outgroup not shown). The ultrafast bootstrap values 99-100% are represented by the asterisks at the nodes. The position of the genus *Laspinema* is highlighted by the arrow.

**FIGURE 2.**
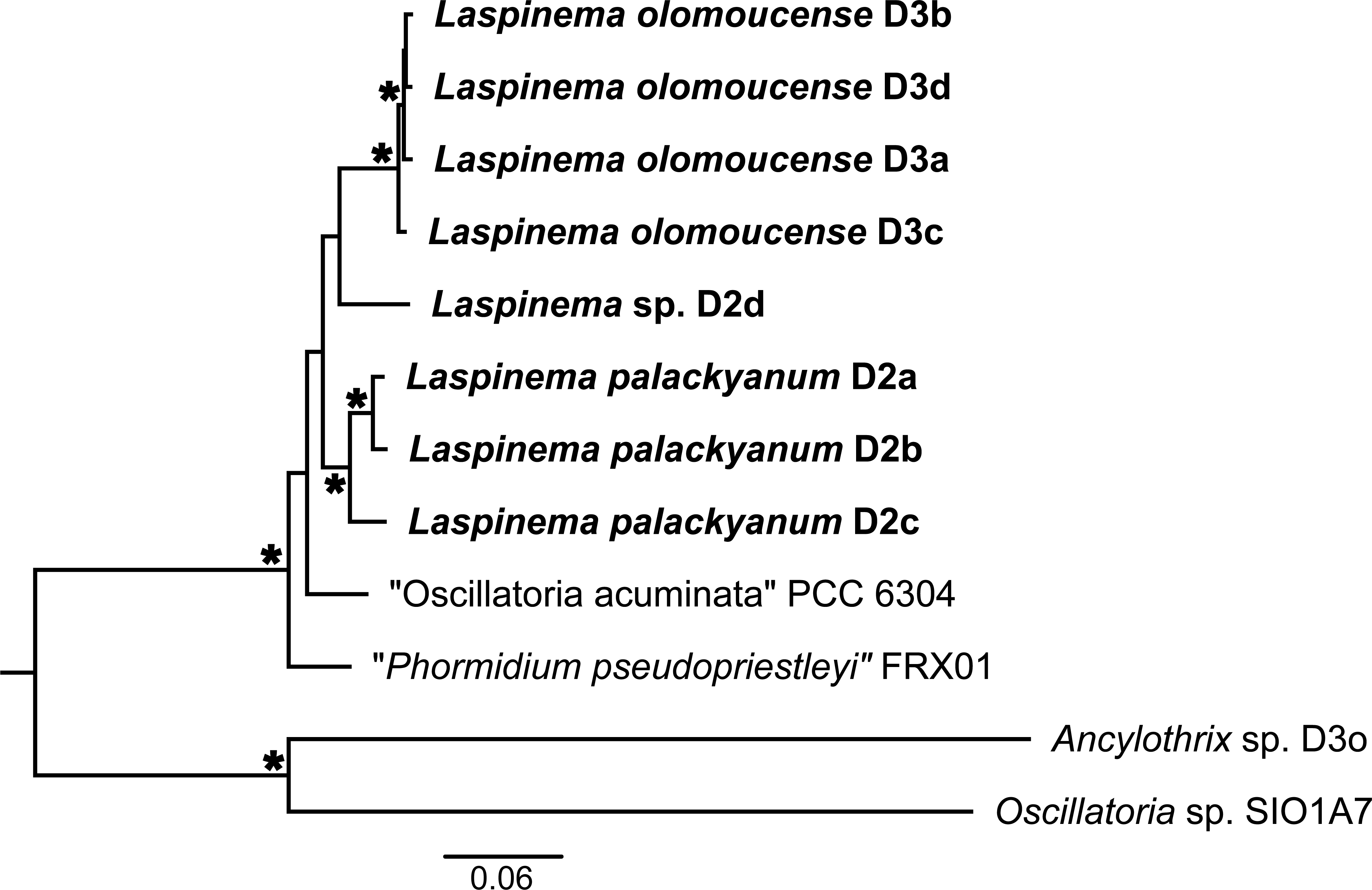
The whole-genome phylogeny of the genus *Laspinema*. The maximum likelihood was reconstructed on the amino acid multiple sequence alignment and rooted to the *Ancylothrix* sp. The ultrafast bootstrap values 99-100% are represented by the asterisks at the nodes. The studied strains are in bold.

The reference strain for the genus *Laspinema* is not available (CCALA culture collection, pers. comm.); therefore, we were unable to sequence its genome. Moreover, 16S rRNA phylogeny fails to recognize the species of *Laspinema* (Stanojković et al. 2022). To resolve this issue, we employed concatenated 16S rRNA and 16S-23S ITS phylogenetic reconstruction. The phylogeny provides evidence that our strains *L. palackyanum* and *L. olomoucense* cluster outside of the reference sequences *L. thermale* proposed in Heidari et al. (2018), although the bootstrap supports were low in the whole tree (Figure 3, Figure S1). We gathered 15 strains from 7 localities across Central Europe, suggesting that *L. olomoucense* represents a widely spread cyanobacterium of the soil habitats. All *L. olomoucense* formed a monophyletic clade. Furthermore, “*Oscillatoria acuminata”* PCC 6304 (with *Lyngbya lagerheimii* SAG 24.99) and *Laspinema* sp. D2d clusters outside of all *Laspinema* species (Figures 2 and 3), which supports that they are separate species of *Laspinema*. The position of “*Phormidium pseudopriestleyi”* FRX01 also suggests that it is one of the *Laspinema* species (Figure 2).

**FIGURE 3.**
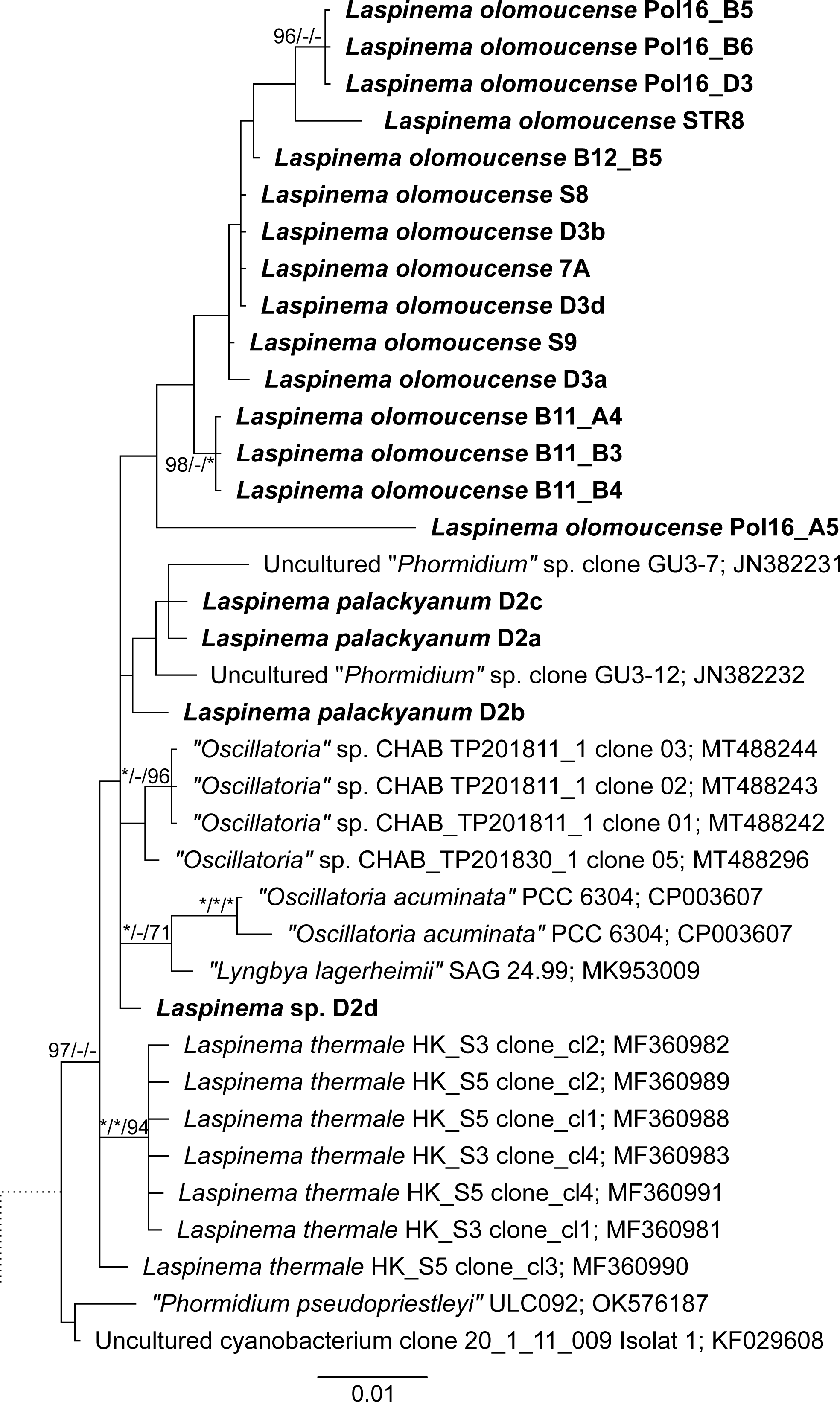
The Bayesian phylogenetic reconstruction based on 16S rRNA and 16S-23S ITS. The tree was rooted to the *Ancylothrix* sp. The investigated strains are in bold. The support values at the nodes are following: BI posterior probability/ML ultrafast bootstrap/MP bootstrap. Values 99-100 are represented by the asterisks at the nodes. Values >70 (MP) and >95 (ML and BI) are shown.

The last phylogenetic tree was reconstructed in order to infer the geographical and ecological patterns within the *Laspinema* at the genus level. *Laspinema* seems to be extremely ecologically versatile with cosmopolitan distribution (Figure 4). *Laspinema* strains were found in soils, mangroves saline or brackish habitats, hot springs, and polar microbial mats in lakes (see a complete list in Table S1). Even expanded 16S rRNA phylogeny fails to recognize the species of *Laspinema* due to the low diversity of the sequences (see below).

**FIGURE 4.**
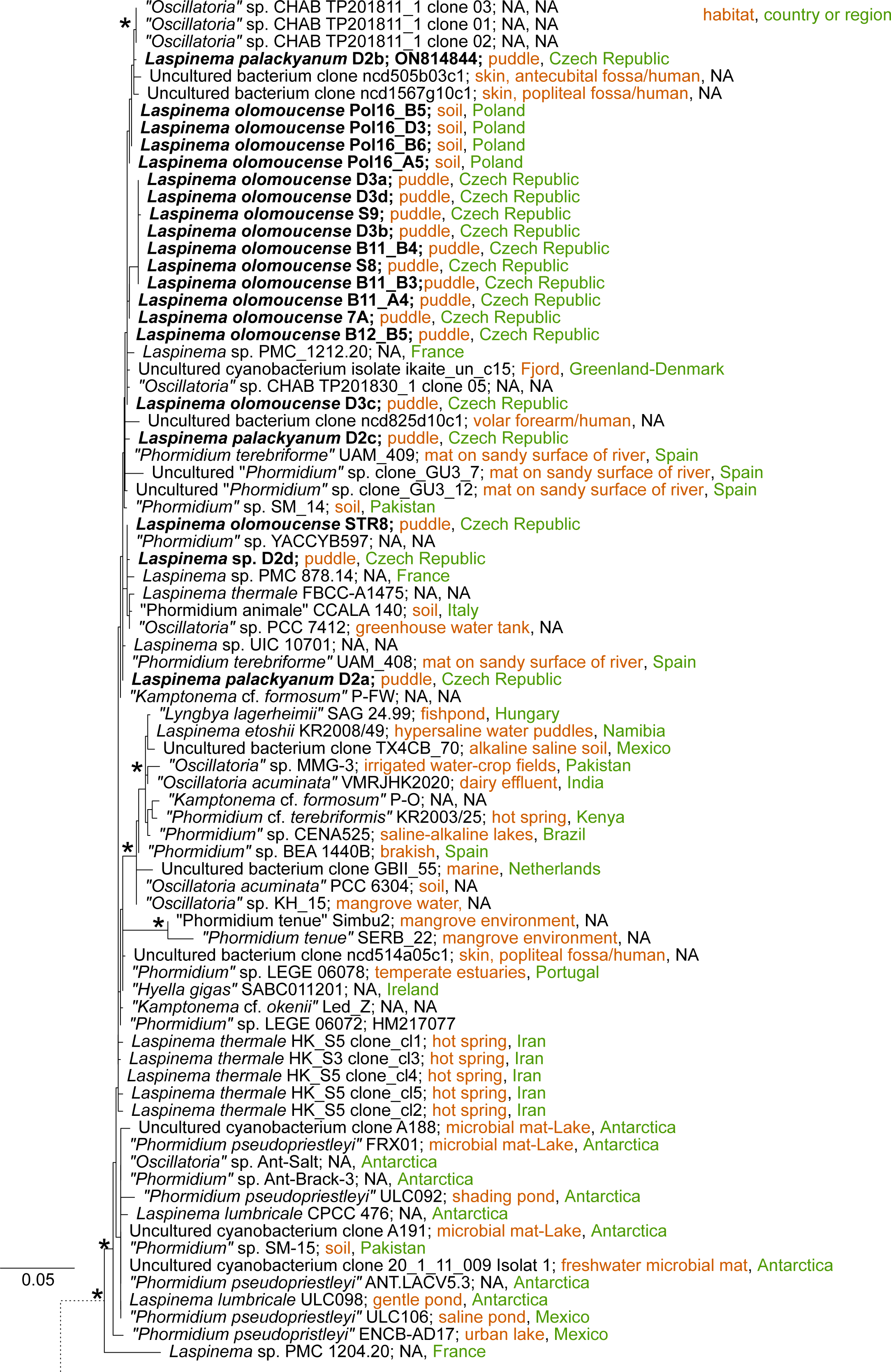
The phylogenetic reconstruction based on 16S rRNA. The maximum likelihood was rooted to the *Gloeobacter violaceus*. The investigated strains are in bold. The ultrafast bootstrap values 99-100% are represented by the asterisks at the nodes. The rest of the phylogenetic tree can be found in supplements (Figure S1).

### Similarity thresholds and ITS secondary structures

Similarity values are often used to define the species boundaries. We performed an estimation of average nucleotide identity (ANI) based on the whole-genome sequences. The 95% ANI cut-off revealed several clusters within *Laspinema* (Table 1). The pairwise comparison of all *L. olomoucense* (D3a-d) strains showed ANI 97-98%. On the other hand, only the species pair *L. palackyanum* D2a and D2b exhibited ANI values over 95%. Strain *L. palackyanum* D2c, *Laspinema* sp. D2d, “*Oscillatoria acuminata”* PCC 6304 and “*Phormidium pseudopriestleyi”* FRX01 had all the ANI values well below the 95% threshold in all pairwise comparisons.

**TABLE 1.**
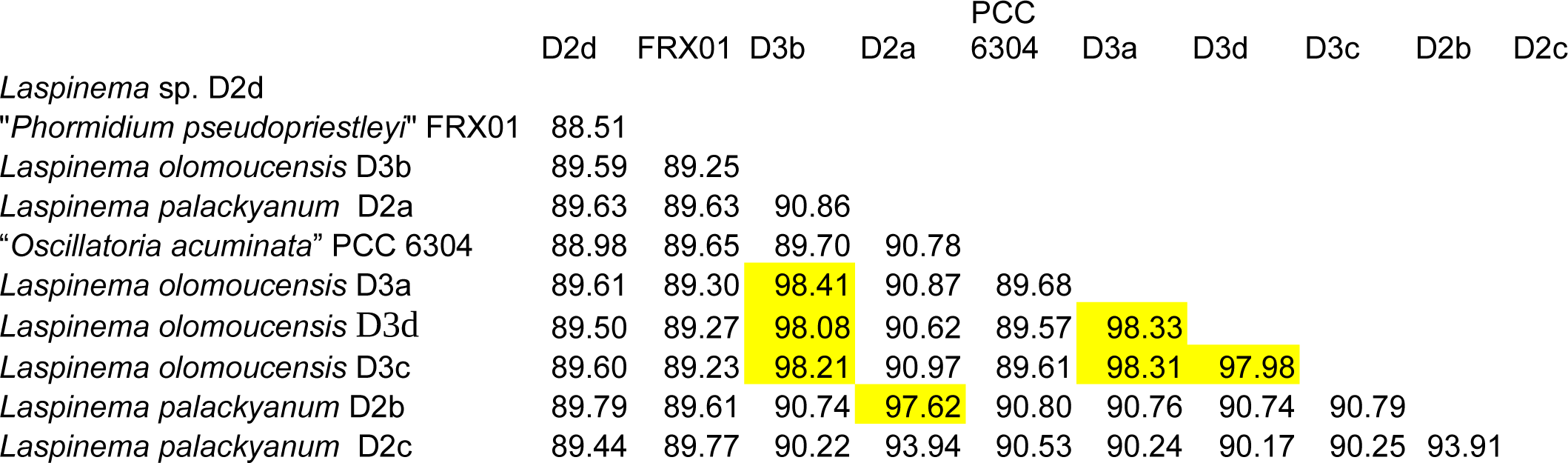
Average nucleotide identities (ANI) of the studied genomes. Threshold >95% values are highlighted in yellow.

We tested two 16S rRNA similarity thresholds within the *Laspinema* genus, 97.5% (Stackebrandt and Goebel 1994) and 98.7% (Stackebrandt and Ebers 2006). We did not identify any specific species clustering, but rather several divergent strains, which revealed similarities below the threshold. Three strains exhibited similarity lower than 97.5% with the rest of the strains (*Phormidium tenue* Simbu2, *Phormidium tenue* SERB-22, and *Laspinema* sp. PMC 1204-20). Several additional strains or metagenomic sequences showed lower similarity than 98.7% with the rest of the strains (Uncultured *Phormidium* sp. clone GU3_7, Uncultured bacterium clone GBII 55, *Kamptonema* cf. *formosum* P_O, *Oscillatoria* sp. MMG-3, Uncultured bacterium clone TX4CB-70). The 16S rRNA similarity did not supply resolution to recognize proposed clusters by the phylogenomic analysis (Table S3).

The secondary structures (D1D1’ and Box B) varied within and between *L. palackyanum* and *L. olomoucense.* We observed nine different D1D1’ and ten different Box B structures (Fig S2 and S3), but their similarity did not resemble the branching of the whole-genome tree (Figure 2) or 16S rRNA and ITS trees (Figure 3). We could not identify specific secondary structure patterns of each species except for the larger terminal bulb of the *L. olomucense,* but this shape was not observed in strains D3a, STR8, B11_B3, and B11_B4.

### Formal description

#### *Laspinema palackyanum* sp. nov. Dvořák et al

##### Description

Filamentous cyanobacterium, colonies macroscopic, growing in mats. The cells of *L. palackyanum* have 4.06 µm mean width (range 3.26–4.76 µm) and 2.19 µm mean length (range 1.38–3.28 µm; Figure 5). Filaments are straight to slightly curved, blue-green to pale gray-green in color. Thin and diffluent sheaths sometimes present. No true branching was observed. Trichomes are cylindrical, attenuated towards the ends. Attenuated trichome apices are curved, attenuation usually involves 3 to 6 cells, but up to 9 or more can be attenuated. Trichomes slightly constricted at cell walls. Cell content often granulated. Reproduction by necridic cells and subsequent breaking of the filaments into hormogonia. The morphological description was based on both culture and fresh material.

**FIGURE 5.**
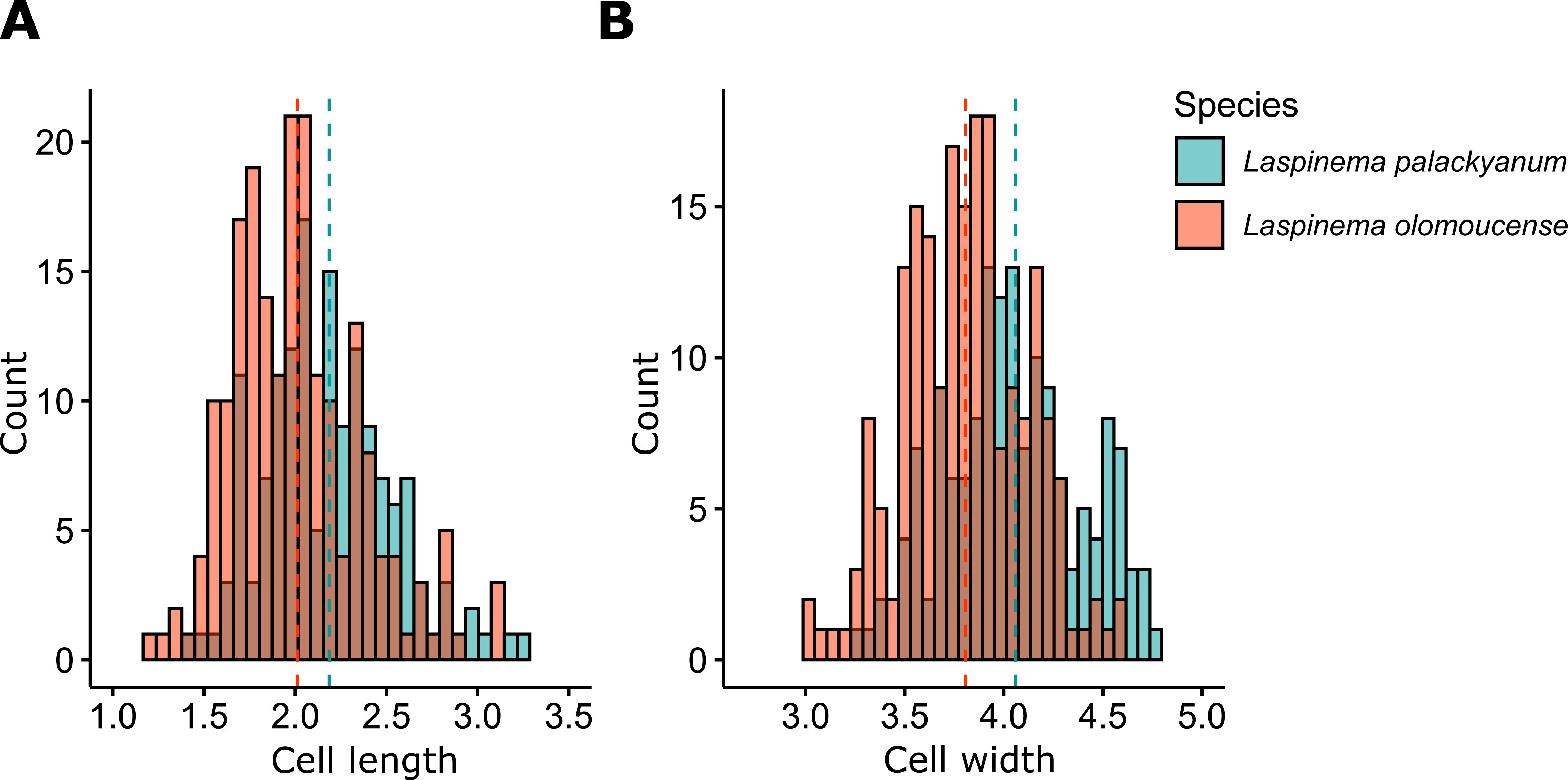
Histograms of cell dimensions of *L. palackyanum* and *L. olomoucense*. Panel A contains lengths and panel B less widths. *L. palackyanum* is represented by green and *L. olomoucense* by orange color. Dashed lines indicate the average values of cell dimensions.

##### Holotype

a dried herbarium specimen was stored at the Herbarium of the Department of Botany (Palacký University Olomouc), ID 100 197,

##### Iconotype

Figures 6 and 7

**FIGURE 6.**
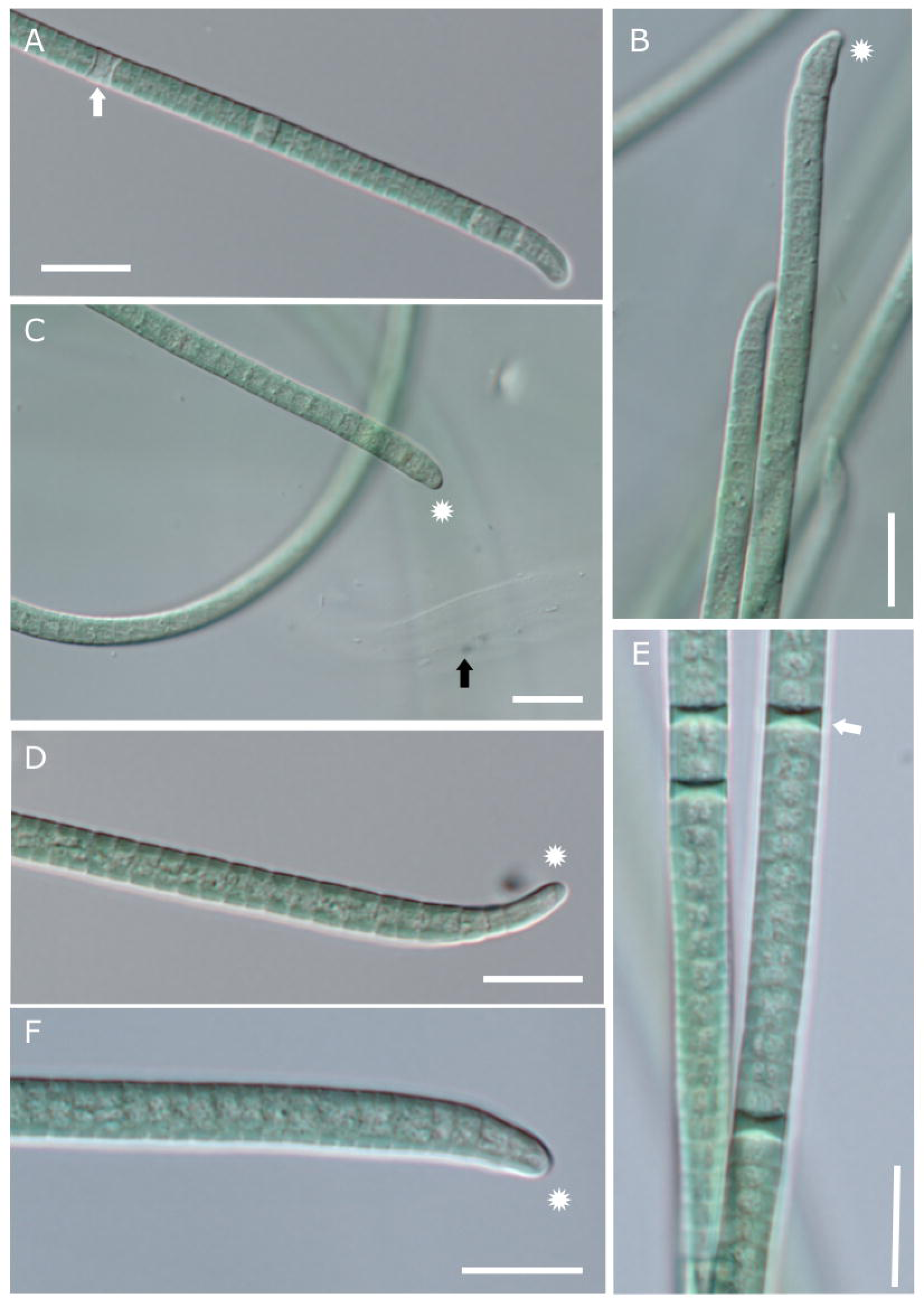
Microphotographs of *Laspinema olomoucense* (A-C) and *Laspinema palackyanum* (D- F). Trichomes of *L. palackyanum* appear to have more pronounced attenuation towards the filament apex (D, marked by asterisk) than in *L. olomoucense* (C, marked by asterisk). This pronounced attenuation is, however, not present in all filaments, as seen in F. Scale = 10 µm, asterisk = attenuated filament apex, wide arrow = necridic cells, black arrow = empty sheath.

**FIGURE 7.**
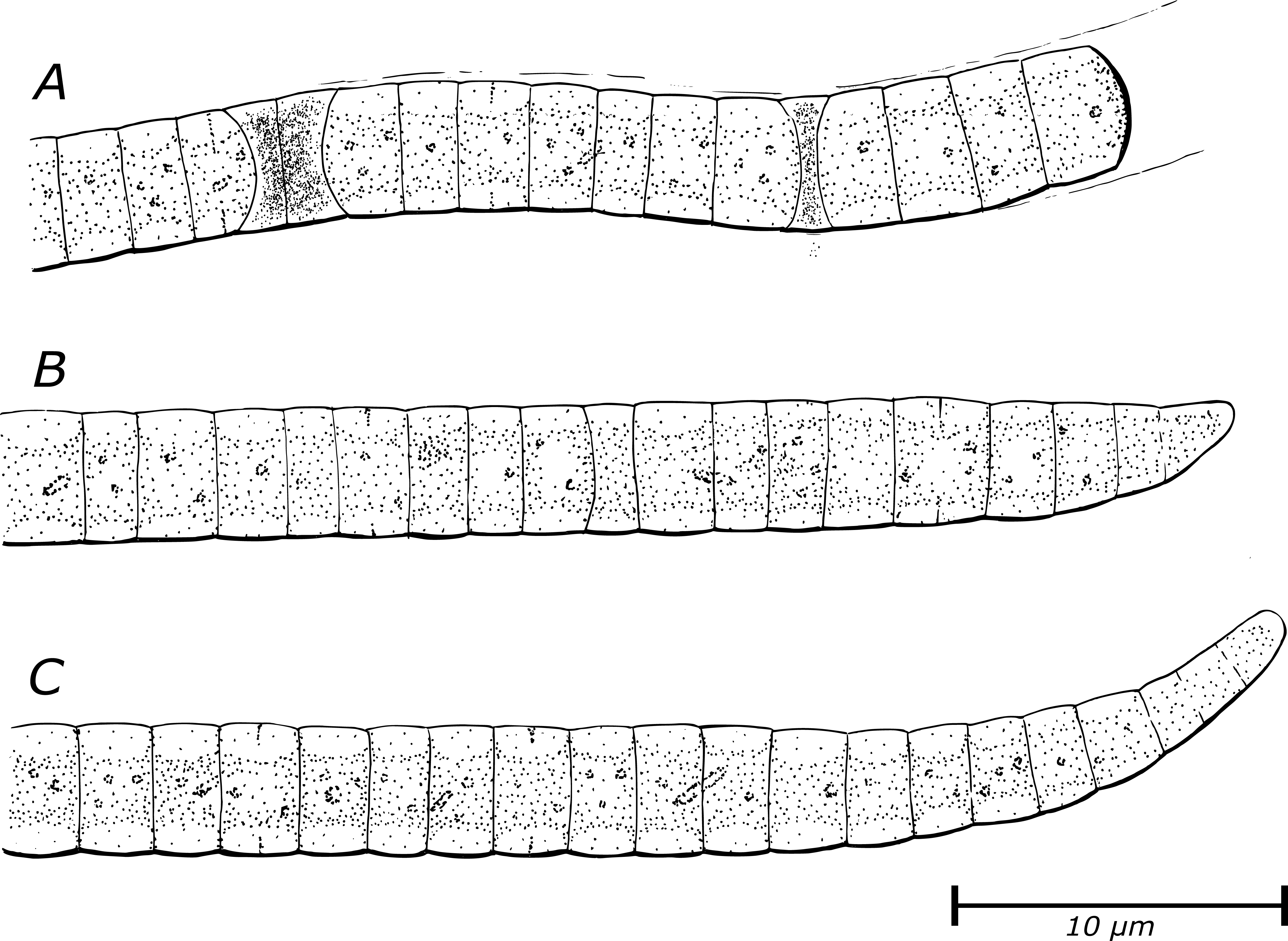
Iconotype of the two new *Laspinema* species. A filament with blunt end and necridic cell (A). A filament of *L. olomoucense* (B) and *L. palackyanym* (C).

**FIGURE 8.**
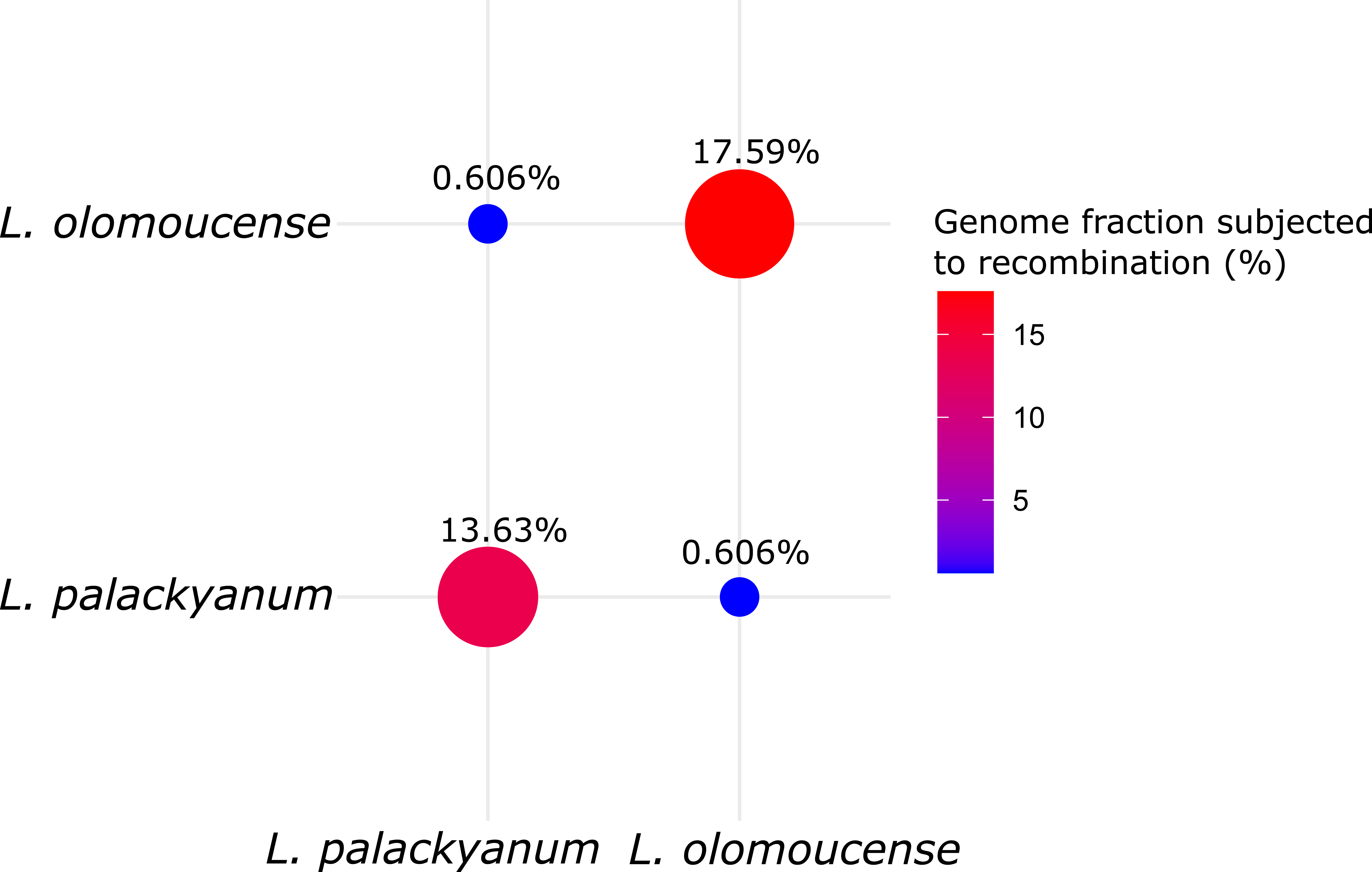
Gene flow bubble plot. Color gradient represents genome fraction subjected to homologous recombination between and within *L. palackyanum* and *L. olomoucense*.

##### Reference strain

*L. palackyanum* D2a stored at CCALA culture collection (Třeboň, Czech Republic). Genbank accession numbers: 16S rRNA (ON814845), ITS (ON814853), and genome assembly (JAMXFF000000000)

##### Habitat

a visible extensive (∼0.5 m^2^) mat covering an ephemeral shallow rainfall pool – puddle.

##### Type locality

a puddle, Olomouc, Czechia (GPS: 49.5766981 N, 17.2653558 E)

##### Etymology

the new species epithet is proposed to honor František Palacký (1798-1876). He was a famous historian and politician and he is often referred to as “Father of the Czech Nation”. Moreover, the epithet celebrates the 450 years anniversary of the Palacký University Olomouc held in 2023.

#### *Laspinema olomoucense* sp. nov. Dvořák et al

##### Description

Filamentous cyanobacterium, colonies macroscopic, growing in mats. The cells of *L. olomoucense* have 3.81 µm mean width (range 3.01–4.58 µm) and 2.01 µm mean length (range 1.23–3.13 µm; Figure 5). Filaments are straight to slightly curved, blue-green to pale gray-green in color. Thin and diffluent sheaths sometimes present. No true branching was observed. Trichomes are cylindrical, attenuated towards the ends. Attenuated trichome apices are curved, attenuation usually involves 3 to 6 cells. Trichomes slightly constricted at cell walls. Cell content often granulated. Reproduction by necridic cells and subsequent breaking of the filaments into hormogonia. The morphological description was based on both culture and fresh material.

##### Holotype

a dried herbarium specimen was stored at the Herbarium of the Department of Botany (Palacký University Olomouc), ID 100 196,

##### Iconotype

Figures 6 and 7

##### Reference strain

1. *L. olomoucense* D3b stored at CCALA culture collection (Třeboň, Czech Republic). Genbank accession numbers: 16S rRNA (ON814840), ITS (ON814848), and genome assembly (JAMXFA000000000)

##### Habitat

a visible extensive (∼0.5 m^2^) mat covering an ephemeral shallow rainfall pool – puddle.

##### Type locality

a puddle, Olomouc, Czechia (GPS: 49.5766981 N, 17.2653558 E)

##### Etymology

the new species epithet is proposed based on the name of the city where we identified the specie for the first time. It honors the ancient city of Olomouc (Czech Republic).

##### Diagnosis

the two new species were derived from *L. thermale* based on the phylogenetic reconstruction. We observed significant difference between *L. palackyanum* and *L. olomoucense* in the cell dimensions (ANOVA, length F = 21.01, width F = 52.78, p < 0.001; Figure 5). However, the cell dimensions largely overlap between the two species, which may prevent precise identification. Some filaments of *L. palackyanum* have up to nine cells or more at the attenuated apex, but this does not seem to be a stable character. Although 16S rRNA phylogeny showed relatively extensive diversity within the genus (Figure 4), it failed to provide a clear clustering pattern as the whole-genome based phylogeny and concatenated phylogeny of 16S rRNA and ITS (Figure 3). The species *L. palackyanum* and *L. olomoucense* could be safely recognized using ANI value and/or phylogenomic analysis. The topology of 16S rRNA and ITS was not significantly supported; therefore, it may not have sufficient resolution to distinguish the two cryptic species. *L. palackyanum* and *L. olomoucense* could be distinguished from *L. lumbricale* and *L. etoshii* because the latter two species have wider cells (Dadheech et al. 2013, Heidari et al. 2018).

## Discussion

Cryptic species currently belong to the most burning issues of the taxonomy of cyanobacteria and other organisms (Osorio-Santos et al. 2014, Stanojković et al. 2022). While most cryptic species could be recognized based on 16S rRNA or ITS secondary structure, *Laspinema olomoucense* and *L. palackyanum* stretch the methodological and technical sensitivity and requirements even further. We need to employ whole-genome sequencing at the population level.

The species definition boils down to a quest to identify of the boundaries between the species. Phylogenetic trees, morphology or other characters could provide measures of difference between the taxa. However, what is enough difference to call the lineages species (Pietrasiak et al. 2019)? To tackle this question, we will adopt the population genomics framework, we recently proposed (Dvořák et al. 2023). We provide several lines of evidence for the evolutionary meaningful species delimitation, albeit the species will remain morphologically cryptic for now. (1) The phylogenomic and phylogenetic inference of 16S rRNA and ITS clearly separated strains of *L. olomoucense* from the strains of *L. palackyanum* (Figures 2 and 3). (2) All pairwise comparisons between *L. palackyanum* and *L. olomoucense* revealed an ANI value way below the threshold of 95% suggested based on the empirical studies (Jain et al. 2018) (Table 1). The ANI value has been criticized by us and other authors because it does not reflect the different rates of speciation among the evolving lineages (Dvořák et al. 2023, Stanojković et al. 2024). However, in the case of *Laspinema*, we cannot compare or estimate rates of speciation and ANI values are relatively low; therefore, we conclude that ANI could safely recognize the species of *Laspinema*. The only exception represents the strain *L. palackyanum* D2c, which showed ANI below the 95% threshold with the rest of the strains. However, its position in the phylogenetic inferences supports that all D2a-c strains belong to the same species *L. palackyanum.* (3) Stanojković et al. (2022) measured overall divergence using the F_ST_ metrics between *L. olomoucense* and *L. palackyanum.* It should be noted that the two species were not proposed in the mentioned paper, but the genomic dataset is the same as analyzed here. F_ST_ (also called fixation index) varies between 0 and 1, where 0 represents mixed populations or species with low divergence and high gene flow. Value 1 represents a complete divergence with no gene flow. The overall F_ST_ value between the two *Laspinema* species was 0.69, suggesting a high divergence because values over 0.25 often indicate a significant divergence between the species (Wright 1965). (4) The pangenome of the two species varied significantly. It is expected that each strain would have its own set of genes as we showed in *Laspinema* (Stanojković et al. 2022). This suggests that a significant difference must occur among the species. (Willis and Woodhouse 2020) attempted to distinguish species of *Raphidiopsis*, *Microcystis* and *Prochlorococcus* based on the difference in pangenome composition. However, it may be difficult to draw a line on what is enough of a difference to distinguish species in *Laspinema* and all other cyanobacteria. More detailed analyses with a larger dataset will help to investigate the importance of the pangenome in taxonomy in the future.

(5) Cyanobacteria, like the rest of prokaryotes, display a copious amount of gene flow via homologous recombination (HR) within and between the species (Bobay and Ochman 2017, Arnold et al. 2021). Prokaryotic HR has a mechanism similar to the eukaryotes except for the directionality; prokaryotic HR is only unidirectional, from donor to recipient (Arnold et al. 2021). In any case, the evolutionary patterns introduced by HR in prokaryotes resemble those in eukaryotes (reviewed in Bobay and Ochman 2017, Kollár et al. 2022). This includes a higher probability of HR between closely related individuals, because the HR molecular machinery can utilize only highly similar DNA fragments. Thus, we could apply the biological species concept (Mayr 1942), which conceptualizes a species as a group of interbreeding organisms sexually isolated from other organisms. This could be modified to prokaryotes using the gene flow in the form of HR – a species is a group of organisms or populations interconnected with gene flow but separated from other organisms by a barrier to gene flow (Bobay and Ochman 2017, Dvořák et al. 2023). Based on this concept, we recognize two species *– L. palackyanum* and *L. olomoucense* because they share only a 0.606% fraction of the genome via HR (Stanojković et al. 2022). Thus, the gene flow between the species is negligible, although the genome fraction subjected to HR within the species was estimated to be 13.63% (*L. palackyanum*) and 17.59% (*L. olomoucense*) (Figure 7).

The 16S rRNA phylogeny failed to recognize the species of *Laspinema.* As shown in many other cyanobacterial taxa, 16S rRNA lacks a taxonomic resolution at the species level reviewed in Dvořák et al. (2023). However, it brought important insights into the ecology and biogeography of the genus *Laspinema*. The strains of *Laspinema* were isolated around the globe from hyper saline to freshwater environments, soils, puddles, water, hot springs, and the human body. This evidences that the cyanobacterial genera are composed of many ecologically diverse species, although many were initially described as monospecific or few species genera. High ecological variability suggests that the number of species in *Laspinema* will increase with a growing number of sequenced genomes. Furthermore, the genetic databases contain a multitude of sources for our understanding of geographical and ecological diversity, which helps to expand our understanding of cyanobacterial distribution and ecology (Skoupý et al. 2022).

We showed that population genomics represents a viable and effective approach to the species delimitation in cyanobacteria. We anticipate that population genomics will grow in significance with the decreasing price of sequencing, development of the bioinformatical tools, and population sampling datasets.

## Acknowledgments

We would like to express our gratitude to Michael D. Guiry (National University of Ireland, Galway, Ireland and AlgaeBase; https://www.algaebase.org/) for help with the species epithets form. The research was funded by the Internal Agency of Palacký University (grant no. IGA PrF-2024- 001).

## Data availability statement

The 16S rRNA and ITS sequence accession numbers are listed in Table S1 and the genome accession numbers are in the Table S2. We stored multiple sequence alignments at figshare (DOI: 10.6084/m9.figshare.24551170) as well as Table S3 (DOI: 10.6084/m9.figshare.24551158).

## Author Contributions

**Petr Dvořák**: conceptualization (lead); funding acquisition (lead); formal analysis (equal); writing – Original Draft Preparation (lead). **Svatopluk Skoupý**: formal analysis (equal); investigation (equal); writing – Review & Editing (equal). **Kateřina Páleníčková**: investigation (equal). **Hana Jarošová**: investigation (equal). **Aleksandar Stanojković**: formal analysis (equal); investigation (lead); writing – Review & Editing (equal).

## Supplementary Materials

**Figure S1.** The phylogenetic reconstruction based on 16S rRNA. The maximum likelihood was rooted to the *Gloeobacter violaceus*. The investigated strains are in bold. The ultrafast bootstrap values 99-100% are represented by the asterisks at the nodes.

**Figure S2.** ITS secondary structures of D1D1’ helices of *Laspinema* strains investigated in this study.

**Figure S3.** ITS secondary structures of Box B helices of *Laspinema* strains investigated in this study.

**Table S1.** A list of strains analyzed in this study and list of 16S rRNA and ITS GenBank accessions with characterization of habitat and locality.

**Table S2.** A list of the GenBank accession numbers of all genomes analyzed in this study.

**Table S3.** The 16S rRNA distance matrix of the *Laspinema* sequences. The values represent p- values inferred by MEGA X.

## References

Arnold, B.J., Huang, I.T. & Hanage, W.P. 2021. Horizontal gene transfer and adaptive evolution in bacteria. Nat. Rev. Microbiol. 2021 204. 20:206–18.

Bobay, L.-M. & Ochman, H. 2017. Biological Species Are Universal across Life’s Domains. Genome Biol. Evol. 9:491–501.

Boyer, S.L., Flechtner, V.R. & Johansen, J.R. 2001. Is the 16S-23S rRNA internal transcribed spacer region a good tool for use in molecular systematics and population genetics? A case study in cyanobacteria. Mol. Biol. Evol. 18:1057–69.

Capella-Gutiérrez, S., Silla-Martínez, J.M. & Gabaldón, T. 2009. trimAl: a tool for automated alignment trimming in large-scale phylogenetic analyses. Bioinformatics. 25:1972–3.

Dadheech, P.K., Casamatta, D.A., Casper, P. & Krienitz, L. 2013. Phormidium etoshii sp. nov. (Oscillatoriales, Cyanobacteria) described from the Etosha Pan, Namibia, based on morphological, molecular and ecological features. Fottea. 13:235–44.

Dvorák, P., Casamatta, D.A., Hašler, P., Jahodárová, E., Norwich, A.R. & Poulícková, A. 2017. Diversity of the cyanobacteria. In Modern Topics in the Phototrophic Prokaryotes: Environmental and Applied Aspects. pp. 3–46.

Dvořák, P., Casamatta, D.A., Poulíčková, A., Hašler, P., Ondřej, V. & Sanges, R. 2014a. *Synechococcus*: 3 billion years of global dominance. Mol. Ecol. 23:5538–51.

Dvořák, P., Hašler, P., Casamatta, D.A. & Poulíčková, A. 2021. Underestimated cyanobacterial diversity: trends and perspectives of research in tropical environments. Fottea. 21:110–27.

Dvořák, P., Hašler, P., Pitelková, P., Tabáková, P., Casamatta, D.A. & Poulíčková, A. 2017. A new cyanobacterium from the everglades, Florida - Chamaethrix gen. nov. Fottea. 17:269–76.

Dvořák, P., Hindák, F., Hašler, P., Hindáková, A. & Poulíčková, A. 2014b. Morphological and molecular studies of *Neosynechococcus sphagnicola*, gen. et sp. nov. (Cyanobacteria, Synechococcales). Phytotaxa. 170:24–34.

Dvořák, P., Jahodářová, E., Stanojković, A., Skoupý, S. & Casamatta, D.A. 2023. Population genomics meets the taxonomy of cyanobacteria. Algal Res. 72:103128.

Dvořák, P., Poulíčková, A., Hašler, P., Belli, M., Casamatta, D.A. & Papini, A. 2015. Species concepts and speciation factors in cyanobacteria, with connection to the problems of diversity and classification. Biodivers. Conserv. 24:739–57.

Edgar, R.C. 2004. MUSCLE: multiple sequence alignment with high accuracy and high throughput. Nucleic Acids Res. 32:1792–7.

Emms, D.M. & Kelly, S. 2015. OrthoFinder: solving fundamental biases in whole genome comparisons dramatically improves orthogroup inference accuracy. Genome Biol. 16:157.

Engene, N., Rottacker, E.C., Kaštovský, J., Byrum, T., Choi, H., Ellisman, M.H., Komárek, J. et al. 2012. *Moorea producens* gen. nov., sp. nov. and Moorea bouillonii comb. nov., tropical marine cyanobacteria rich in bioactive secondary metabolites. Int. J. Syst. Evol. Microbiol. 62:1171–8.

Flombaum, P., Gallegos, J.L., Gordillo, R.A., Rincón, J., Zabala, L.L., Jiao, N., Karl, D.M. et al. 2013. Present and future global distributions of the marine Cyanobacteria *Prochlorococcus* and *Synechococcus*. Proc. Natl. Acad. Sci. U. S. A. 110:9824–9.

Fox, G.E., Wisotzkey, J.D. & Jurtshuk, P. 1992. How close is close: 16S rRNA sequence identity may not be sufficient to guarantee species identity. Int. J. Syst. Bacteriol. 42:166–70.

Hassler, H.B., Probert, B., Moore, C., Lawson, E., Jackson, R.W., Russell, B.T. & Richards, V.P. 2022. Phylogenies of the 16S rRNA gene and its hypervariable regions lack concordance with core genome phylogenies. Microbiome 2022 101. 10:1–18.

Heidari, F., Zima, J., Riahi, H. & Hauer, T. 2018. New simple trichal cyanobacterial taxa isolated from radioactive thermal springs. Fottea. 18:137–49.

Hoang, D.T., Chernomor, O., Von Haeseler, A., Minh, B.Q. & Vinh, L.S. 2018. UFBoot2: Improving the Ultrafast Bootstrap Approximation. Mol. Biol. Evol. 35:518–22.

Jahodářová, E., Dvořák, P., Hašler, P., Holušová, K. & Poulíčková, A. 2018. *Elainella* gen. nov.: a new tropical cyanobacterium characterized using a complex genomic approach. Eur. J. Phycol. 53:39–51.

Jain, C., Rodriguez-R, L.M., Phillippy, A.M., Konstantinidis, K.T. & Aluru, S. 2018. High throughput ANI analysis of 90K prokaryotic genomes reveals clear species boundaries. Nat. Commun. 2018 *91*. 9:1–8.

Johansen, J.R. & Casamatta, D.A. 2005. Recognizing cyanobacterial diversity through adoption of a new species paradigm. Algol. Stud. für Hydrobiol. Suppl. Vol. 117:71–93.

Johansen, J.R., Mareš, J., Pietrasiak, N., Bohunická, M., Zima, J., Štenclová, L. & Hauer, T. 2017. Highly divergent 16S rRNA sequences in ribosomal operons of *Scytonema hyalinum* (Cyanobacteria). PLoS One. 12:e0186393.

Kalyaanamoorthy, S., Minh, B.Q., Wong, T.K.F., Von Haeseler, A. & Jermiin, L.S. 2017. ModelFinder: fast model selection for accurate phylogenetic estimates. Nat. Methods 2017 146. 14:587–9.

Kollár, J., Poulíčková, A. & Dvořák, P. 2022. On the relativity of species, or the probabilistic solution to the species problem. Mol. Ecol. 31:411–8.

Komárek, J., Johansen, J.R., Šmarda, J. & Strunecký, O. 2020. Phylogeny and taxonomy of *Synechococcus*-like cyanobacteria. Fottea. 20:171–91.

Komárek, J., Kaštovský, J., Mareš, J. & Johansen, J.R. 2014. Taxonomic classification of cyanoprokaryotes (cyanobacterial genera) 2014, using a polyphasic approach. Preslia. 86:295– 335.

Kumar, S., Stecher, G., Li, M., Knyaz, C. & Tamura, K. 2018. MEGA X: Molecular Evolutionary Genetics Analysis across Computing Platforms. Mol. Biol. Evol. 35:1547–9.

Labrada, N.A., McGovern, C.A., Thomas, A.L., Hurley, A.C., Mooney, M.R. & Casamatta, D.A. 2023. The CIMS (Cyanobacterial ITS motif slicer) for molecular systematics. Fottea.

Larsson, A. 2014. AliView: a fast and lightweight alignment viewer and editor for large datasets. Bioinformatics. 30:3276–8.

Mayr, E. 1942. Systematics and the Origin of Species. Columbia Univ. Press, New York.

Nabout, J.C., da Silva Rocha, B., Carneiro, F.M. & Sant’Anna, C.L. 2013. How many species of Cyanobacteria are there? Using a discovery curve to predict the species number. Biodivers. Conserv. 22:2907–18.

Nguyen, L.-T., Schmidt, H.A., von Haeseler, A. & Minh, B.Q. 2015. IQ-TREE: a fast and effective stochastic algorithm for estimating maximum-likelihood phylogenies. Mol. Biol. Evol. 32:268– 74.

Osorio-Santos, K., Pietrasiak, N., Bohunická, M., Miscoe, L.H., Kováčik, L., Martin, M.P. & Johansen, J.R. 2014. Seven new species of *Oculatella* (Pseudanabaenales, Cyanobacteria): taxonomically recognizing cryptic diversification. Eur. J. Phycol. 49:450–70.

Pietrasiak, N., Osorio-Santos, K., Shalygin, S., Martin, M.P. & Johansen, J.R. 2019. When Is A Lineage A Species? A Case Study In Myxacorys gen. nov. (Synechococcales: Cyanobacteria) With The Description of Two New Species From The Americas. J. Phycol. 55:976–96.

R Core Team 2021. A Language and Environment for Statistical Computing. *R Found. Stat. Comput. Vienna*, Austria.

Ronquist, F., Teslenko, M., Van Der Mark, P., Ayres, D.L., Darling, A., Höhna, S., Larget, B. et al. 2012. MrBayes 3.2: Efficient Bayesian Phylogenetic Inference and Model Choice Across a Large Model Space. Syst. Biol. 61:539.

Skoupý, S., Stanojković, A., Casamatta, D.A., McGovern, C.A., Martinović, A., Konderlová, M., Dodoková, V. et al. 2024. Population genomics and morphological data bridge the centuries of cyanobacterial taxonomy along the continuum of *Microcoleus* species. iScience.

Skoupý, S., Stanojković, A., Pavlíková, M., Poulíčková, A. & Dvořák, P. 2022. New cyanobacterial genus *Argonema* is hiding in soil crusts around the world. Sci. Reports 2022 121. 12:1–15.

Stackebrandt, E. & Ebers, J. 2006. Taxonomic parameters revisited: Tarnished gold standards. Microbiol. Today 33:152–5.

Stackebrandt, E. & Goebel, B.M. 1994. Taxonomic note: A place for DNA-DNA reassociation and 16S rRNA sequence analysis in the present species definition in bacteriology. Int. J. Syst. Bacteriol. 44:846–9.

Stanojković, A., Skoupý, S., Johannesson, H. & Dvořák, P. 2024. The global speciation continuum of the cyanobacterium *Microcoleus*. Nat. Commun.

Stanojković, A., Skoupý, S., Škaloud, P. & Dvořák, P. 2022. High genomic differentiation and limited gene flow indicate recent cryptic speciation within the genus Laspinema (cyanobacteria). Front. Microbiol. 13:977454.

Turner, S., Pryer, K.M., Miao, V.P.W. & Palmer, J.D. 1999. Investigating Deep Phylogenetic Relationships among Cyanobacteria and Plastids by Small Subunit rRNA Sequence Analysis1. J. Eukaryot. Microbiol. 46:327–38.

Willis, A. & Woodhouse, J.N. 2020. Defining Cyanobacterial Species: Diversity and Description Through Genomics. Crit. Rev. Plant Sci. 39:101–24.

Wright, S. 1965. The Interpretation of Population Structure by F-Statistics with Special Regard to Systems of Mating. Evolution (N. Y). 19:395–420.

Zimba, P. V., Shalygin, S., Huang, I.S., Momčilović, M. & Abdulla, H. 2021. A new boring toxin producer–*Perforafilum tunnelli* gen. & sp. nov. (Oscillatoriales, Cyanobacteria) isolated from Laguna Madre, Texas, USA. Phycologia. 60:10–24.

Zuker, M. 2003. Mfold web server for nucleic acid folding and hybridization prediction. Nucleic Acids Res. 31:3406–15.

