## Supplementary figures and images for "Population genomics resolves cryptic species of the ecologically flexible genus *Laspinema* (cyanobacteria)^1^"

### Fig. S1

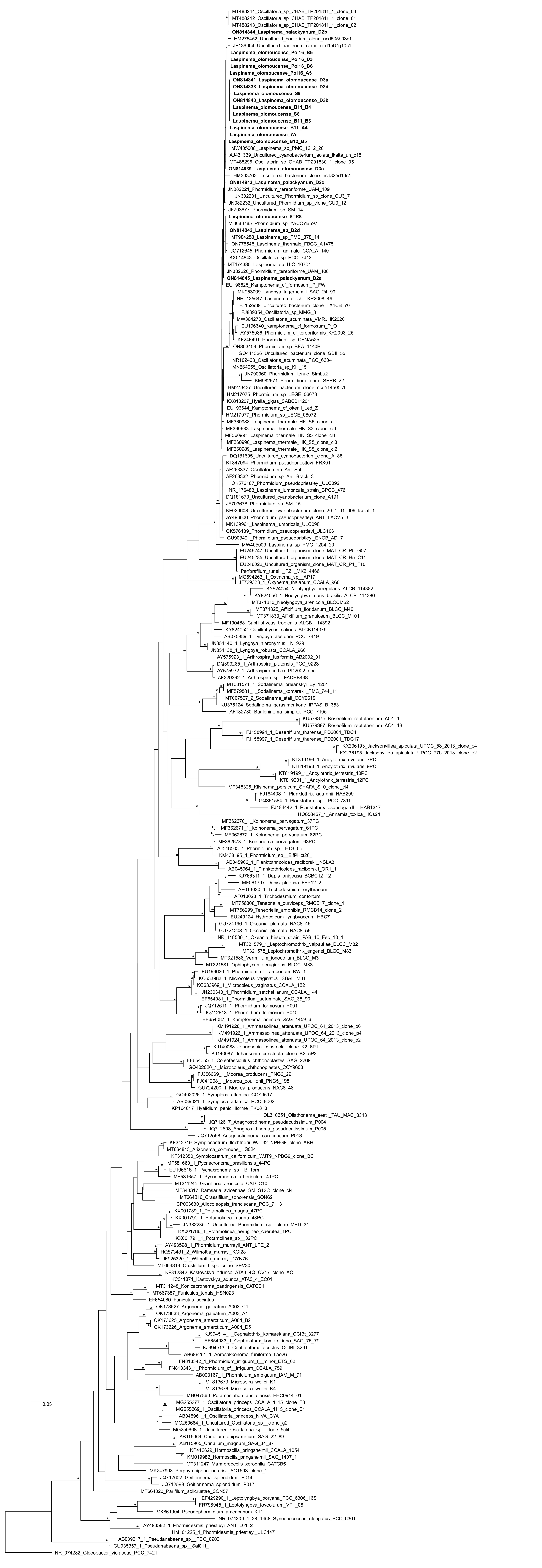
