## Supplementary material for "Population genomics resolves cryptic species of the ecologically flexible genus *Laspinema* (cyanobacteria)^1^": Fig. S2

D1D1'

*Laspinema palackyanum*

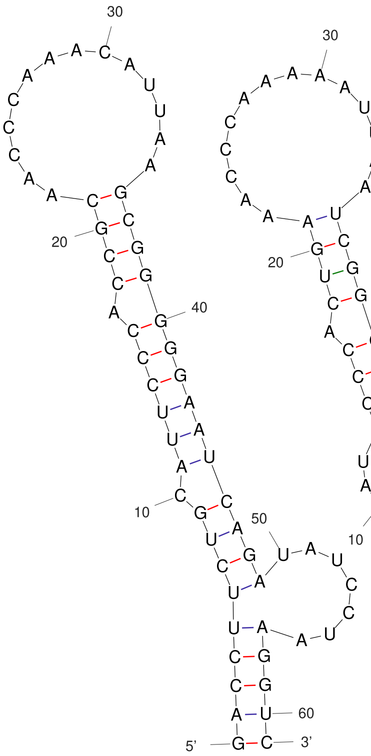

D2a
D2c

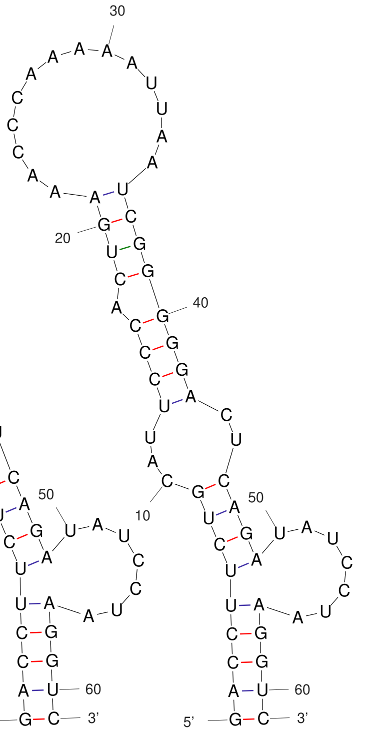

D2b

*Laspinema* sp.

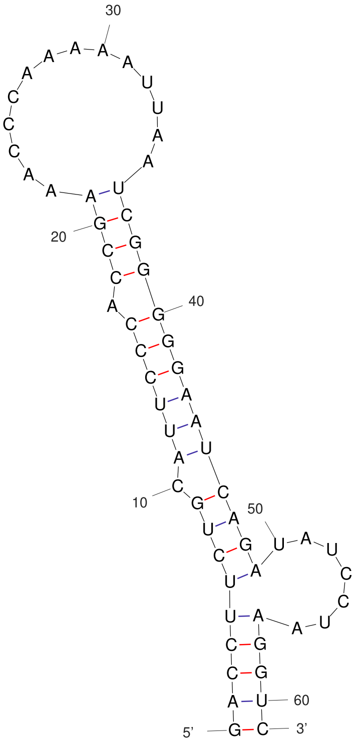

D2d

*Laspinema olomoucense*

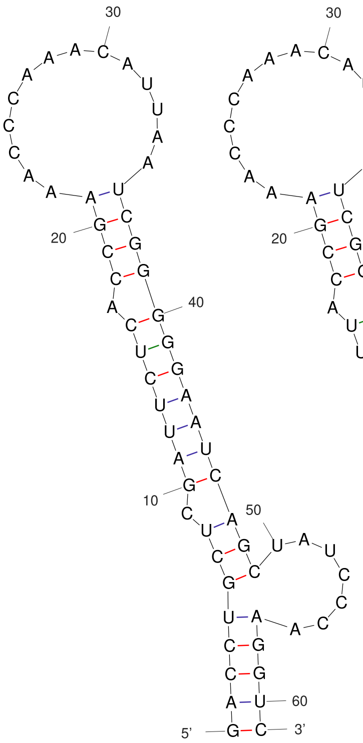

D3a
D3b
D3d
S8

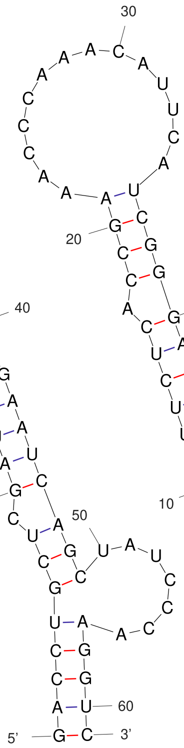

Pol16_B5
Pol16_B6
Pol16_D3

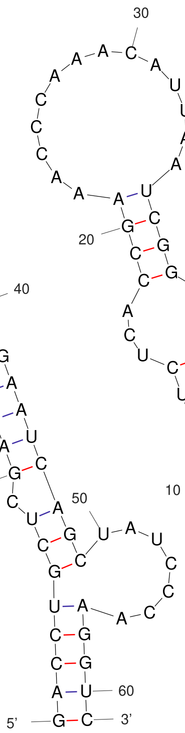

STR8

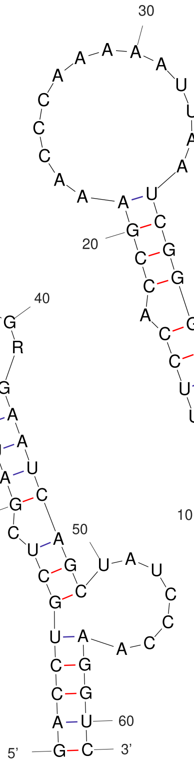

7A

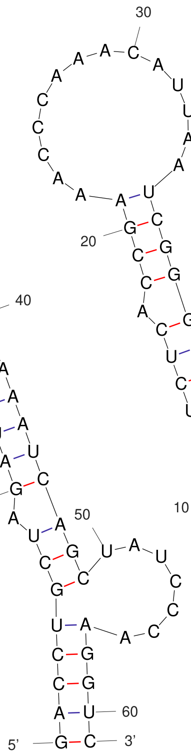

B11_A4
B11_B3
B11_B4

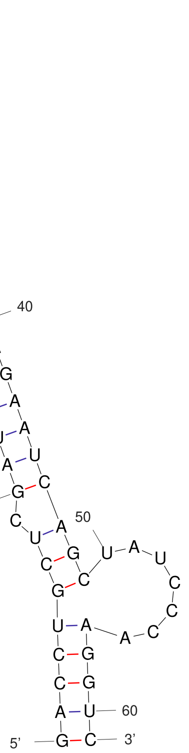

B12_B5
