## Supplementary material for "Population genomics resolves cryptic species of the ecologically flexible genus *Laspinema* (cyanobacteria)^1^": Fig. S3

Box B

*Laspinema palackyanum*

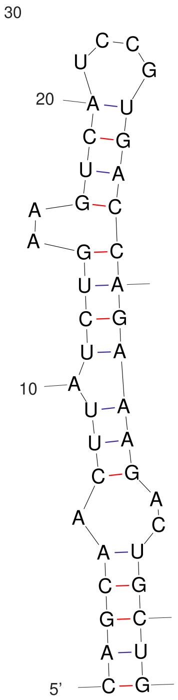

D2a
D2c

*Laspinema* sp.

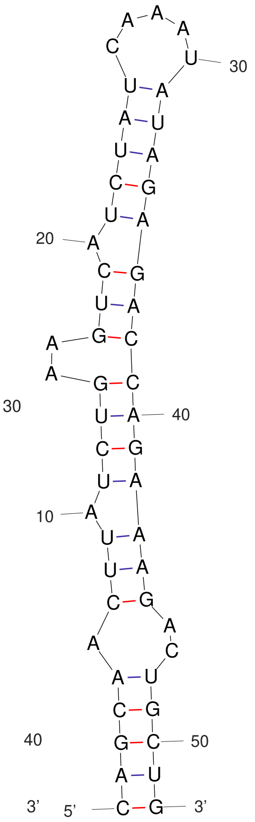

D2d

*Laspinema olomoucense*

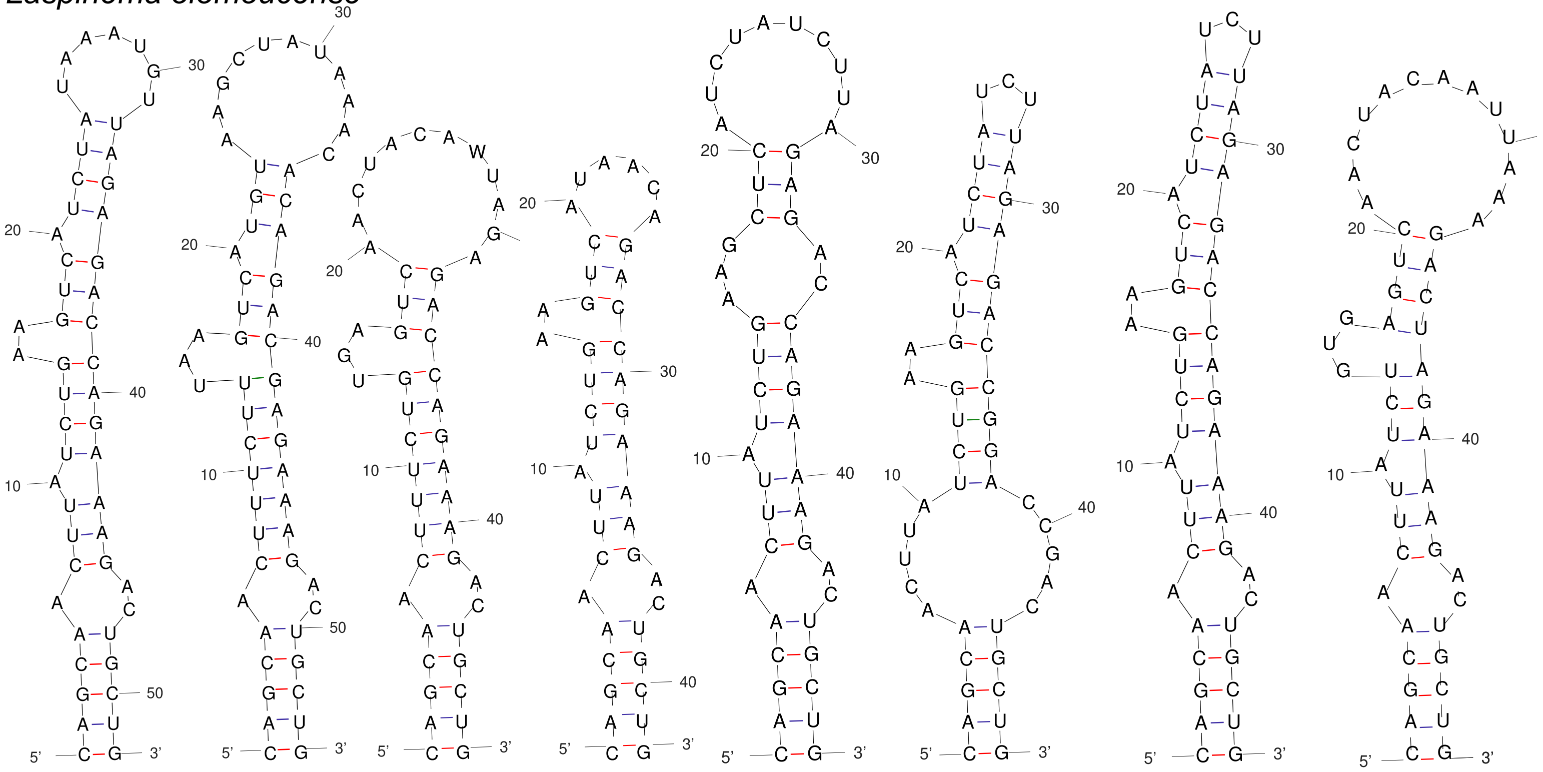

D3a | Pol16_B5, Pol16_B6, Pol16_D3 | S8 | STR8 | B11_A4 | B11_B3 | B11_B4 | B11_B5
